# Biological Mechanisms of Strength Preservation During Calorie Restriction-Induced Weight Loss Among Young- to Middle-Aged Adults without Obesity

**DOI:** 10.64898/2026.01.29.702632

**Authors:** Nadia Moore, Akshay Bareja, Leanna M. Ross, Katherine A. Collins-Bennett, Susan B. Racette, Leanne M. Redman, Corby K. Martin, Sai Krupa Das, William E. Kraus, Kim M. Huffman

## Abstract

**Purpose:** Weight loss is often pursued to improve cardiometabolic health and quality of life. However, rapid weight loss can lead to reductions in lean soft tissue mass and strength to compromise body composition and functional ability. Thus, identifying molecular predictors of muscular strength preservation during weight loss is critical to mitigating these effects.

**Methods:** We conducted a secondary analysis of the CALERIE^TM^ (Comprehensive Assessment of Long-term Effects of Reducing Intake of Energy) trial, a two-year randomized controlled study of caloric restriction (CR) or *ad-libitum* intake in healthy adults without obesity. Among 198 participants, changes in whole-body mass and knee extensor strength were assessed over the first 12 months of the study which was primarily characterized by weight loss. Transcriptomic profiling was conducted in a subset of 42 participants who provided skeletal muscle samples. Linear regression was used to model the relationship between strength change and gene expression change, while controlling for changes in whole-body mass. Gene set enrichment analysis (GSEA) was performed using Hallmark pathways. Individual-level pathway analysis was performed via gene set variation analysis (GSVA).

**Results:** We identified 96 out of 198 individuals (48.5%) who maintained or improved strength relative to body mass during weight loss (*i.e.* individuals with residuals > 0). Transcriptomics analysis on a subset of 42 individuals revealed 151 genes significantly associated with change in strength after accounting for change in whole-body mass (*p* < 0.01). Hub genes were identified as *HSP90AA1* (β = 34.45, SE = 7.32, *p* <0.001), *EIF3A* (β = 36.14, SE = 10.27, *p* < 0.001), *EIF5B* (β = 49.94, SE = 11.28, *p* < 0.001), and *H3C1* (β = −15.87, SE = 4.85, *p* < 0.001). GSEA revealed significant involvement of pathways related to cellular proliferation, immune regulation, protein secretion, and checkpoint control processes. GSVA identified a similar set of pathways.

**Conclusions:** These findings highlight molecular pathways supporting strength retention during CR-induced weight loss. Heat-shock protein, *HSP90AA1*, warrants further investigation as a candidate target for preserving muscle strength during interventions aimed at weight reduction.

## INTRODUCTION

Approximately 42% of adults worldwide are actively attempting to lose weight.^1^ Weight loss is often undertaken to improve cardiometabolic disease risk factors, including insulin resistance, blood pressure, and lipid profiles.^2^ However, weight loss can also be associated with adverse health consequences, particularly in older adults whose body composition is comprised of more adipose relative to lean mass. On average, 14% to 23% of weight lost is lean mass,^3,4^ with greater degrees of weight loss—particularly losses exceeding 10% of baseline body weight—being associated with reductions in lean mass and bone mineral density.^3,5^ When these reductions are substantial, they may contribute to negative health outcomes, including decreased basal metabolic rate, compromised bone health, increased fracture risk, and reduced muscle strength.^6–11^ While weight loss can improve mobility for many individuals with obesity, excessive losses of lean mass and bone mineral density may offset these gains in other individuals. Given the well-documented risk of muscle and strength loss during weight reduction^12^, there is a growing need to identify molecular predictors of strength retention and gain during weight loss.

As an initial step toward this goal, we leveraged physiologic and skeletal muscle transcriptomic data from the Comprehensive Assessment of Long-term Effects of Reducing Intake of Energy (CALERIE™) 2 trial, which investigated the effects of two years of calorie restriction (CR) on various physiological outcomes among younger to middle-aged (21–50 years) healthy adults without obesity (BMI 22.0–27.9 kg/m²). In CALERIE™, participants undergoing CR for two years experienced reductions in whole-body mass, lean mass (whole-body and leg), cardiorespiratory fitness, and absolute strength. However, when strength was expressed relative to body mass (*i.e.,* relative strength), the CR group on average demonstrated preservation or improvement in relative strength at two years.^11^ These findings highlight that CR-induced weight loss can lead to both strength losses and gains, depending on individual factors. Thus, using gene expression data derived from skeletal muscle tissue, CALERIE™ provides a unique opportunity to identify molecular mediators of strength maintenance and gain during weight loss.

## MATERIALS AND METHODS

### Study Participants

This secondary analysis utilized data from 198 participants of the original 220 individuals enrolled in CALERIE™ Phase 2 – a multicenter, randomized controlled trial conducted from May 2007 to November 2012 (NCT00427193). CALERIE™ Phase 2 was designed to evaluate severe biological effects of 25% CR across two years among healthy men (aged 21-50 years) and premenopausal women (aged 21-47 years) with normal weight or mild to moderate overweight (BMI 22.0-27.9 kg/m^2^).^13^ Participants were randomized in a 2:1 ratio to either the CR behavioral intervention or to the *ad libitum* (AL, non-restricted control) group. The study protocol was approved by the institutional review boards at all participating clinical centers (Washington University School of Medicine, St Louis, MO; Pennington Biomedical Research Center, Baton Rouge, LA; Tufts University, Boston, MA) and the coordinating center (Duke University, Durham, NC). All study participants provided written informed consent, and study oversight was provided by a Data and Safety Monitoring Board. Extensive details of the study design, recruitment and screening process, and intervention are provided in previous publications.^13,14^

### Study Design

At baseline, participants in the CR group were prescribed a 25% reduction in calorie intake based on individualized energy requirements determined from two consecutive doubly labeled water (DLW) measurements conducted over four weeks. For the DLW protocol, urine samples were collected before and at multiple time points following ingestion of water labeled with deuterium (^2^H) and oxygen-18 (^18^O); isotopic enrichment was quantified by isotope ratio mass spectrometry. Total daily energy expenditure (kcal/day) was calculated from the differential elimination rates of the two isotopes.^15^

All participants underwent DLW measurements and dual-energy x-ray absorptiometry (DXA) scans at six-month intervals throughout the two-year intervention. Percent CR was calculated retrospectively using DLW-derived total daily energy expenditure in combination with DXA-derived changes in body composition. As reported previously, due to variability in adherence and limitations of the original weight loss trajectory models used to guide prescriptions, participants in the CR group achieved an average 11.9% CR over two years.^10,16^ Adherence was monitored using a computerized tracking system that allowed real-time adjustments to counseling and support strategies based on individual weight trajectories. Participants assigned to the AL control group were instructed to maintain their habitual diets and received no dietary counseling.

### Study Outcomes

Data collection occurred over a two-year period at multiple time points. As previous findings from CALERIE™ indicated that the first year of intervention was primarily characterized by weight loss and the second year by weight maintenance^10,11,17^, this secondary analysis focused specifically on whole-body mass, muscular strength, and skeletal muscle gene expression data collected at baseline and 12 months. Given the objective of this work was to investigate the biological basis of strength maintenance and gains in the context of weight loss, this analysis did not focus on comparing CR and AL groups. Instead, the analysis evaluated concomitant changes in whole-body mass, muscle strength, and associated skeletal muscle gene expression across participants. Accordingly, study group assignment (CR or AL) was not included as a covariate in subsequent regression models.

#### Anthropometrics

Body weight was measured at baseline and 12 months to the nearest 0.1 kg using a calibrated digital scale, with participants wearing light clothing and no shoes.^10^

#### Muscular strength

Muscle strength of knee extensors and knee flexors were evaluated using Biodex System 3 dynamometers (Biodex Medical Systems, Shirley, NY) at all three clinical centers; a Cybex II Isokinetic Dynamometer (Cybex Division of Lumex, Inc., Ronkonkoma, NY) was used for a subset of participants at the Pennington Biomedical Research Center. Participants performed isokinetic strength tests which included five repetitions of concentric knee extension and concentric knee flexion at 60°/s, then at 180°/s, with a 30-s rest between sets. After a 5-min rest, muscular endurance was assessed using 30 repetitions of concentric knee extension and flexion at 180°/s. After another 5-min rests, participants performed three 5-s maximal isometric contractions at 45° for knee extension and flexion, with a 30 s rest between. The entire sequence was performed on each leg.

#### Skeletal muscle gene expression

At both baseline and 12 months, a subset of participants (n=91) volunteered for optional skeletal muscle biopsies of the vastus lateralis collected using a percutaneous needle biopsy technique under local anesthesia, following standard procedures.^11^ Approximately 100 mg of muscle tissue was obtained, immediately flash-frozen in liquid nitrogen, and stored at –80°C until analysis. Total RNA was extracted using TRIzol reagent (Invitrogen, Carlsbad, CA). RNA integrity was assessed using the Agilent 2100 Bioanalyzer (Agilent Technologies, Santa Clara, CA). mRNA sequencing, filtering, alignment, and genome annotation were performed as described in Das et al. (2023).^18^

### Software and Analytical Tools

All initial data processing and linear regression analyses were conducted using Python libraries: pandas v1.4.4, numpy v1.26.4, matplotlib v3.5.2, seaborn v0.11.2, statsmodels v0.13.2, and scipy v1.9.1.^19–24^ Residuals from these regression models were extracted and used for downstream analyses and data visualizations performed in R (version 4.5.0) and RStudio (version 2025.05.0+496). Additional preprocessing, regression analyses, pathway enrichment, and plotting were also conducted in R. The R libraries used include: dplyr v1.1.4^25^, tidyr v1.3.1^26^, readr v2.1.5^27^, tidyverse v2.0.0^28^, biomaRt v2.60.1^29,30^, edgeR v4.2.2^31–34^, DT v0.33^35^, fgsea v1.30.0^36^, GSVA v2.2.0^37,38^, ggtext v0.1.2^38^, patchwork v1.3.0^39^, pathview v1.44.0^40^, clusterProfiler v4.12.6^41–43^, and org.Hs.eg.db v3.19.1.^44^

### Statistical Analysis

#### Linear Regression

Using Python, a linear regression (*sm.OLS (y, X).fit())* was run on the percent change in strength (y) and percent change in whole-body mass (X) within the baseline to 12-month timeframe **(Eq. 1)**. The residuals from this regression analysis (n = 198) were plotted to ensure normal distribution.

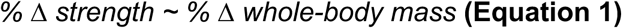

#### Multiple Regression

Raw RNA-seq count data from skeletal muscle samples were imported into R. The raw RNA-seq count data (paired baseline and 12-month) were available for 42 of the 198 participants. The original gene counts matrix had 60,605 genes. Genes were removed if >50% participants had raw counts of zero at either the baseline or 12-month timepoint, resulting in 32,291 remaining genes. This filtering approach is consistent with our earlier study.^18^ The count data were then normalized using the Trimmed Mean of M-values (TMM) method, which were subsequently log_2_ transformed to produce log2-counts-per-million (CPM) data.^45,46^ Of these genes, only genes with corresponding gene symbols were considered in downstream analyses (25,200 remaining genes). Changes in gene expression were calculated as:

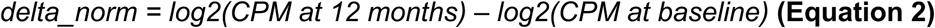

To assess the relationship between change in gene expression and change in strength while accounting for change in mass, (n_genes_ = 25,200; n_participants_ = 42), change in strength was regressed onto change in gene expression and change in mass using **Equation 3**. This regression was performed independently for each gene. *p*-values from the models were adjusted for multiple testing using the Benjamini-Hochberg method. All results were stored as tables.

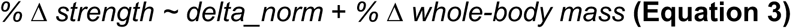

#### Ranking Genes

To prepare for gene set enrichment analysis (GSEA), two types of ranking metrics were calculated. One metric used a composite score of sign (log2 fold change) multiplied by the negative log10-transformed *p*-value for the *delta_norm* covariate in **Equation 3**. This metric incorporated both directionality and statistical confidence. In addition, we used the *delta_norm t*-statistic for each gene. This metric captured both the magnitude and certainty of the association and is a widely accepted gene-ranking approach.^47^ In cases where multiple ensembl gene IDs mapped to the same gene symbol, we used the mean of signed fold-change or *t*-statistic to rank the gene. In the following steps and results, we focus on the results provided by the *t*-statistic gene-ranking method.

#### Gene Set Enrichment Analysis (GSEA)

For pathway analysis, gene set collections were downloaded from the Molecular Signatures Database (MSigDB v2024.1, Broad Institute). Two collections were downloaded from the GSEA website: the Hallmark gene sets and the full MSigDB collection.^48^ Both gene sets were imported using the gmtPathways function. GSEA was performed using the fgseaMultilevel function from the fgsea R package, with default parameters. The output was sorted by nominal *p*-value to prioritize statistically significant pathway associations.

The Hallmark collection results, with *t*-statistics used as the ranking metric, were selected as the primary focus for downstream interpretation. This choice was motivated by the clarity and reduced redundancy of Hallmark gene sets, as well as the consistent interpretability of *t*-statistics across genes. These results were output as a table and were observed for pathways most associated with change in strength after accounting for whole-body mass change. Finally, the top five upregulated and top five downregulated pathways, based on their normalized enrichment scores (NES), were selected as the top ten pathways of interest. The pathways were visualized with their respective NES.

#### Gene Set Variation Analysis (GSVA)

For individual-level pathway analysis, we performed GSVA using the same gene sets as described in the GSEA section. Baseline and 12-month pathway scores were computed for each individual using the *GSVA::gsvaParam()* function from the GSVA R package. log2(CPM) data were used as input and a gaussian kernel was used for the non-parametric estimation of the empirical cumulative distribution function of expression levels across samples. Changes in pathway activity for each individual were computed by subtracting the baseline GSVA score from the 12-month GSVA score for each pathway. In cases where multiple ensembl gene IDs mapped to the same gene symbol, we used the ensembl ID associated with the highest variance across all samples as the “representative”.

#### Visualization and Integration of Genes of Interest

To identify genes associated with strength preservation, we performed the regression described in **Equation 3** for each gene. Protein-coding genes with a nominal *p*-value < 0.01 were retained, and the top five upregulated and top five downregulated genes, ranked by *t*-statistic, were selected as the ten genes of interest (GOIs). Because STRING is limited to protein-coding genes, this subset was used for network analysis. The primary goal of this analysis was to identify potential drug targets by detecting hub genes—highly connected nodes within protein-protein interaction networks. STRING integrates evidence from scientific literature and experimental data to visualize protein-protein interactions. The ten GOIs were submitted to STRING to construct an interaction network for further analysis.

## RESULTS

CALERIE™ participants having both strength and mass data (whom we refer to as the “Overall cohort”) (n = 198) were used to compute residuals. These residuals were then used to visually identify individuals with unusually high or low changes in strength given their changes in whole-body mass (relative to the cohort). Residuals were correlated with GSVA scores to identify pathways of interest. We refer to the participants who also had skeletal muscle transcriptional data as the “Gene expression subset” (n = 42). Characteristics of these two cohorts were recorded (**Table 1**; **Figure 1**). For the Overall cohort, percent changes in strength and whole-body mass were significantly related, but with a weak relationship (*p*<0.001, R² = 0.07) (**Figure 2**). Though the regression showed residuals with a normal distribution, there was a single outlier around 80 Nm (**Figure 3**).

**Figure 1.**
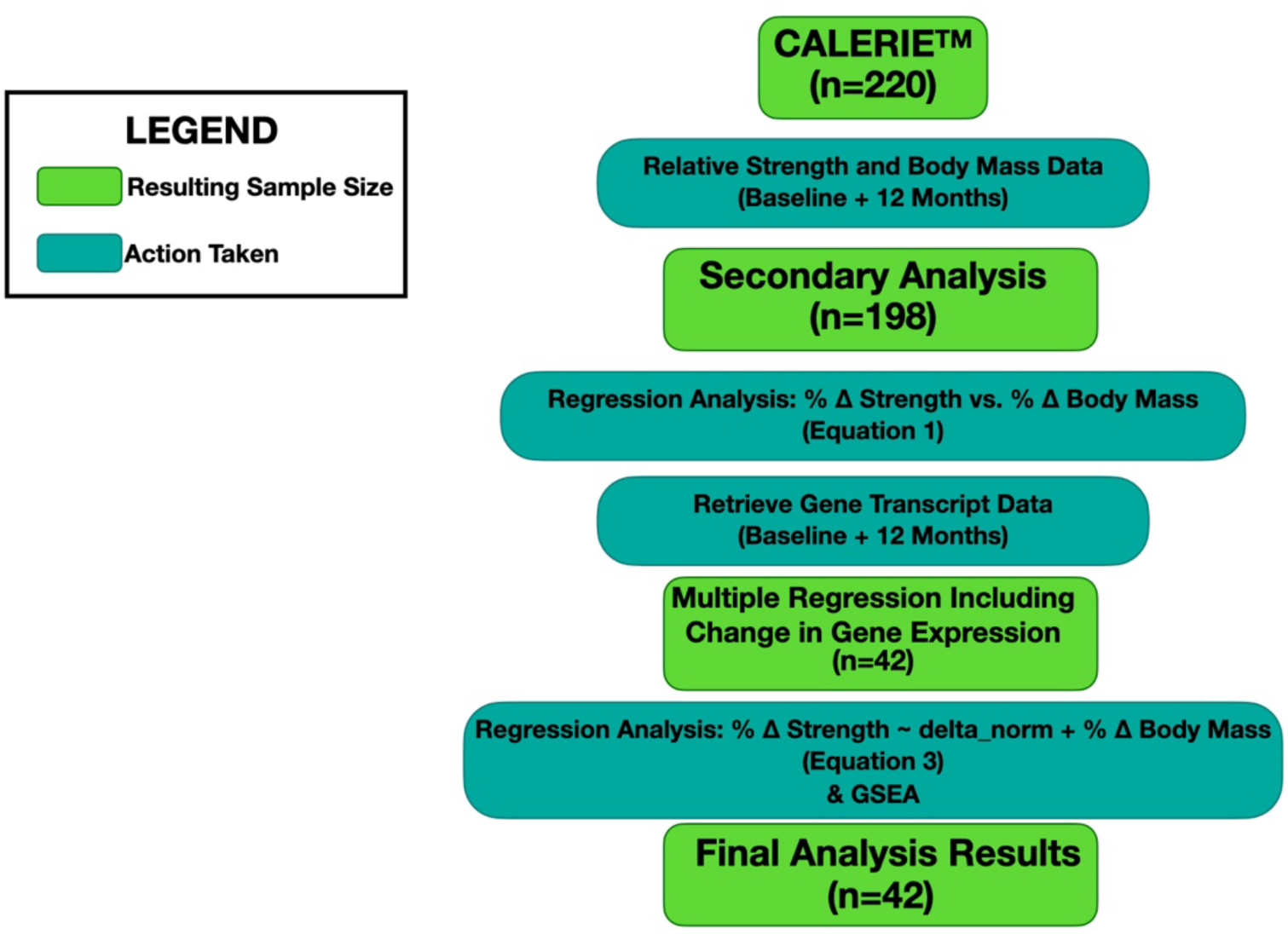
Flowchart for CALERIE Participants with Strength, Mass, and Gene Expression Data. CALERIE™ Phase 2 had 220 participants, 198 of which had strength and whole-body mass measurements at both baseline and 12 months. Changes in strength and body mass were analyzed in a linear regression model that provided residuals (n = 198). Of these 198 individuals, 42 had gene transcript data available at both the baseline and 12-month time points. Multiple linear regression was performed for each gene independently (see **Equation 3** in main text). The results of these regressions were then put through a gene set enrichment analysis. Thus, the final analysis results on genes and gene pathways are provided for only these 42 individuals.

**Figure 2.**
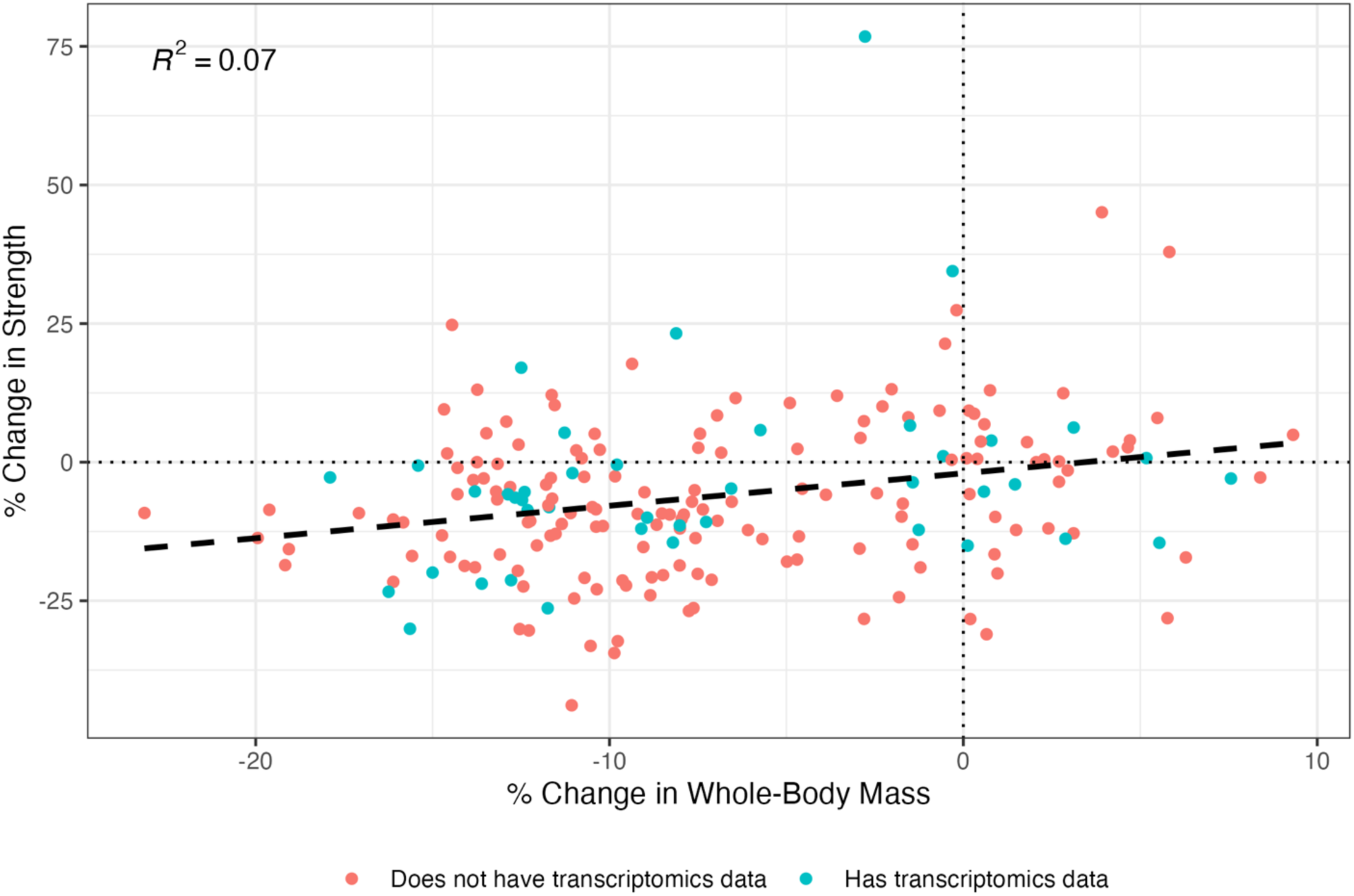
Regression of Percent Change in Strength Against Percent Change in Whole-Body Mass (N=198). Percent change in isometric knee extension strength was regressed against percent change in whole-body mass from baseline to 12 months. A positive trend was observed (p < 0.0001), though the low R² value indicates a weak overall association. Point color indicates whether the individual had corresponding transcriptomics data.

**Figure 3.**
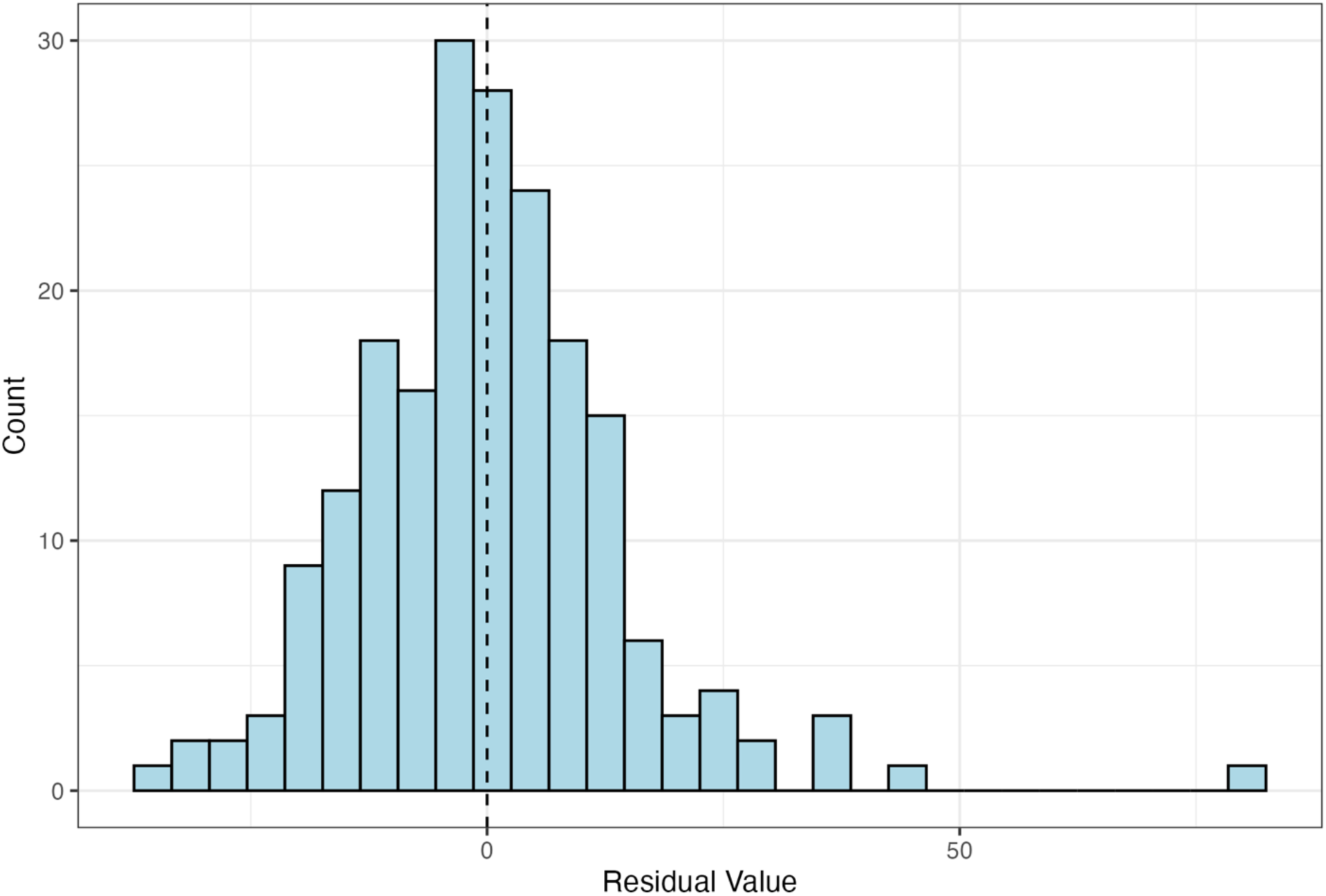
Distribution of Residuals from Strength vs. Whole-Body Mass Regression (N=198). The histogram shows the distribution of residuals from the regression in Figure 2. Residuals were approximately normally distributed, with one high positive outlier (residual ≈ 80).

**Table 1.** Participant characteristics at baseline and 12 months.

| Variables | Overall Cohort (n=198) | Gene Expression Subset (n=42) |
| --- | --- | --- |
| Age, yr | 38.15 (7.17) | 38.87 (6.64) |
| Baseline BMI, kg/m<sup>2</sup> | 25.15 (1.68) | 25.12 (1.62) |
| Month 12 BMI, kg/m<sup>2</sup> | 23.2 (2.17) | 23.08 (2.07) |
| Female, n (%) | 139 (70.2%) | 26 (61.9%) |
| Male, n (%) | 59 (29.8%) | 16 (38.1%) |
| Percent Change in Strength | -6.08 (14.77) | -3.54 (17.75) |
| Percent Change in Whole-Body Mass, kg | -6.97 (6.67) | -6.99 (6.95) |

Summary characteristics for the Overall cohort (n = 198) and the subgroup of those individuals that also had their gene transcript data available (n = 42). Age, BMI at baseline and 12 months, sex distribution, and mean percent changes in strength and whole-body mass are reported as mean (SD).

The smaller Gene expression subset was a representative subset of the full cohort, selected based on the availability of gene expression data. This subset was similar to the Overall cohort in age, BMI, sex distribution, and percent change in their whole-body mass. However, the two cohorts’ percent changes in strength were different, with the Overall cohort having a lower mean (mean = −6.1%; SD = 14.8%) change in strength than that of the Gene expression subset (mean = −3.5%; SD = 17.7%).

In regressions, gene counts (from the Gene expression subset) were used to predict change in strength while accounting for change in whole-body mass. After Benjamini-Hochberg correction, no genes were found to be statistically significant. After using a nominal *p*-value threshold of *p* < 0.01, 151 genes were associated with change in strength while accounting for change in whole-body mass. The top five up-and downregulated genes are shown in **Table 2**. Importantly, we conducted a series of sensitivity analyses that included replacing change in whole-body mass with change in fat-free mass (ffm) or change in lean soft-tissue mass (computed as the difference between ffm and total bone mineral content) in the multiple regression models. Additionally, we refit the model described in Equation 3 using data from the subset of individuals who had lost whole-body mass (N=33). **Supplementary Figure 1** shows that in these three analyses the regression results were largely unchanged.

**Table 2.** Top 5 up- and downregulated genes.

| Gene Name | Beta | Standard Error | t-statistic | p-value |
| --- | --- | --- | --- | --- |
| HSP90AA1 | 34.57 | 7.32 | 4.72 | 3.0197E-05 |
| EIF5B | 49.94 | 11.28 | 4.43 | 7.5193E-05 |
| MT-RNR1 | 17.66 | 4.04 | 4.37 | 8.9201E-05 |
| ZNF451-AS1 | -21.58 | 5.3 | -4.07 | 0.00022133 |
| MT-RNR2 | 18.24 | 4.5 | 4.06 | 0.00023131 |
| MIR3935 | 11.17 | 2.77 | 4.03 | 0.00024817 |
| TMEM184C-DT | -11.69 | 3.08 | -3.8 | 0.00049967 |
| ODC1-DT | -22.34 | 6.05 | -3.69 | 0.00068419 |
| ADGRE4P | -10.82 | 3 | -3.61 | 0.00086505 |
| HNRNPH3 | 35.47 | 10.03 | 3.54 | 0.00106252 |

Genes significantly associated with strength changes independent of mass changes were identified using TMM-normalized RNA-seq data. Linear models were ranked by t-statistic, and the top genes were ordered by nominal p-value. Benjamini-Hochberg adjusted p-values are not shown as they were all approximately 1.

To better understand the potential biological relevance of the genes in our gene expression dataset, we performed GSEA and assessed enrichment of the 50 Hallmark gene sets. Among these gene sets, the most enriched pathways were those related to cellular proliferation (*e.g.,* MYC targets, E2F targets), immune and inflammatory responses (*e.g.,* Allograft Rejection, Inflammatory Response, TNFA Signaling via NFKB), and checkpoint regulation (*e.g.,* G2M Checkpoint) (**Table 3**).

**Table 3.** Top 5 up-and downregulated Hallmark pathways.

| Hallmark Pathway | p-value | Adjusted p-value | Log<sub>2</sub> Error | ES | NES | Size |
| --- | --- | --- | --- | --- | --- | --- |
| ALLOGRAFT REJECTION | 4.0231E-05 | 0.00020116 | 0.56 | -0.35 | -1.73 | 189 |
| IL6 JAK STAT3 SIGNALING | 0.00107366 | 0.00412944 | 0.46 | -0.39 | -1.71 | 82 |
| INFLAMMATORY RESPONSE | 0.00324135 | 0.0108045 | 0.43 | -0.3 | -1.47 | 192 |
| TNFA SIGNALING VIA NFKB | 0.00864989 | 0.02703092 | 0.38 | -0.28 | -1.41 | 192 |
| KRAS SIGNALING UP | 0.03599414 | 0.09998371 | 0.32 | -0.26 | -1.29 | 186 |
| MYC TARGETS V1 | 1.2655E-26 | 6.3275E-25 | 1.34 | 0.59 | 2.93 | 200 |
| G2M CHECKPOINT | 2.46E-11 | 6.1501E-10 | 0.86 | 0.45 | 2.25 | 198 |
| PROTEIN SECRETION | 5.3998E-11 | 8.9997E-10 | 0.85 | 0.56 | 2.47 | 96 |
| E2F TARGETS | 7.2486E-08 | 9.0607E-07 | 0.7 | 0.39 | 1.95 | 200 |
| MTORC1 SIGNALING | 8.0685E-07 | 8.0685E-06 | 0.66 | 0.37 | 1.86 | 200 |

Gene set enrichment analysis (GSEA) was conducted on TMM-normalized gene expression data using Hallmark gene sets ( $n = 50$ ). Pathways were ranked based on $t$ -statistic-derived gene ranks and adjusted $p$ -values. The top 5 most significant downregulated and upregulated pathways are shown.

To further assess the pathways most associated with strength change independent of mass change, the ten pathways with the most pronounced directional associations—comprised of the five pathways with the highest positive NES values and the five with the most negative NES values—were selected for visualization (**Figure 4**). Among these, pathways involved in cellular proliferation and muscle adaptation (MYC Targets, E2F Targets, G2M Checkpoint), immune and inflammatory processes (Allograft Rejection, Inflammatory Response, TNFA Signaling via NFKB), and protein secretion were most strongly represented. To note, two metabolic pathways – oxidative phosphorylation and fatty acid metabolism – were associated with statistically significant positive NES values.

**Figure 4.**
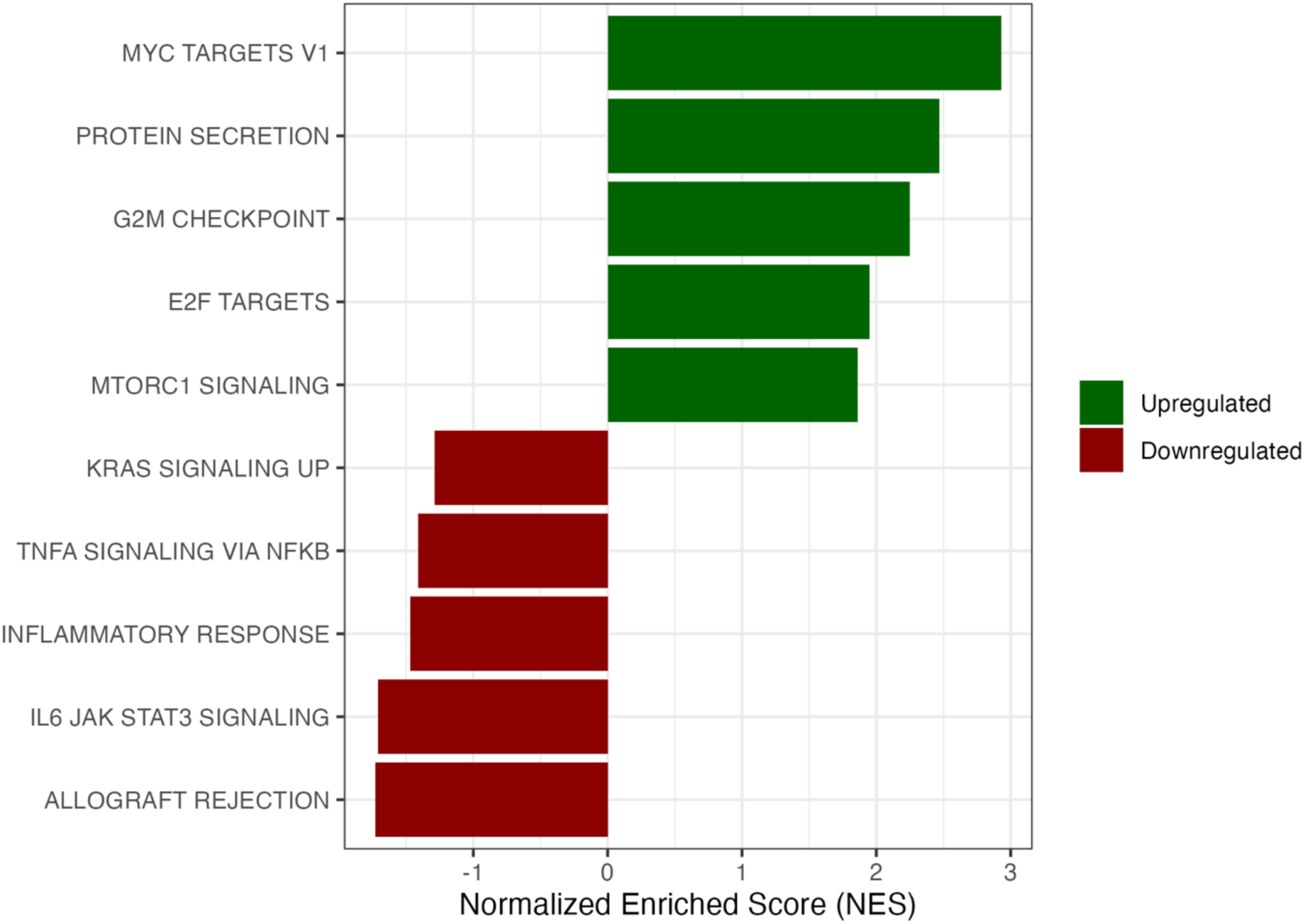
Top 10 Hallmark Pathways Associated with Strength Preservation. The top five positively and top five negatively enriched Hallmark pathways were selected based on absolute magnitude of NES.

To explore gene-level results, the five most upregulated and five most downregulated protein-coding genes were extracted based on their *t*-statistic and a nominal *p*-value of p < 0.01 (**Table 4**). The gene with the most significant positive *t*-statistic, *HSP90AA1*, has prior evidence or plausible functional relevance in the context of muscle maintenance.^49–51^ The top ten protein-coding genes were subsequently analyzed using the STRING database to identify potential functional interactions and intermediary connections. Of the original ten genes, six (*HSP90AA1*, *PF4V1*, *EIF3A, EIF5B*, *H3C1*, and *RAD21*) met the minimum confidence score threshold of 0.900 and demonstrated connections to at least one other gene in the network (**Figure 5**). Notably, *HSP90AA1, EIF3A, EIF5B*, and *H3C1* were the only genes from the initial set indirectly linked to each other within the STRING interaction network. *HSP90AA1* was also the top leading edge gene for the fatty acid metabolism pathway.

**Figure 5.**
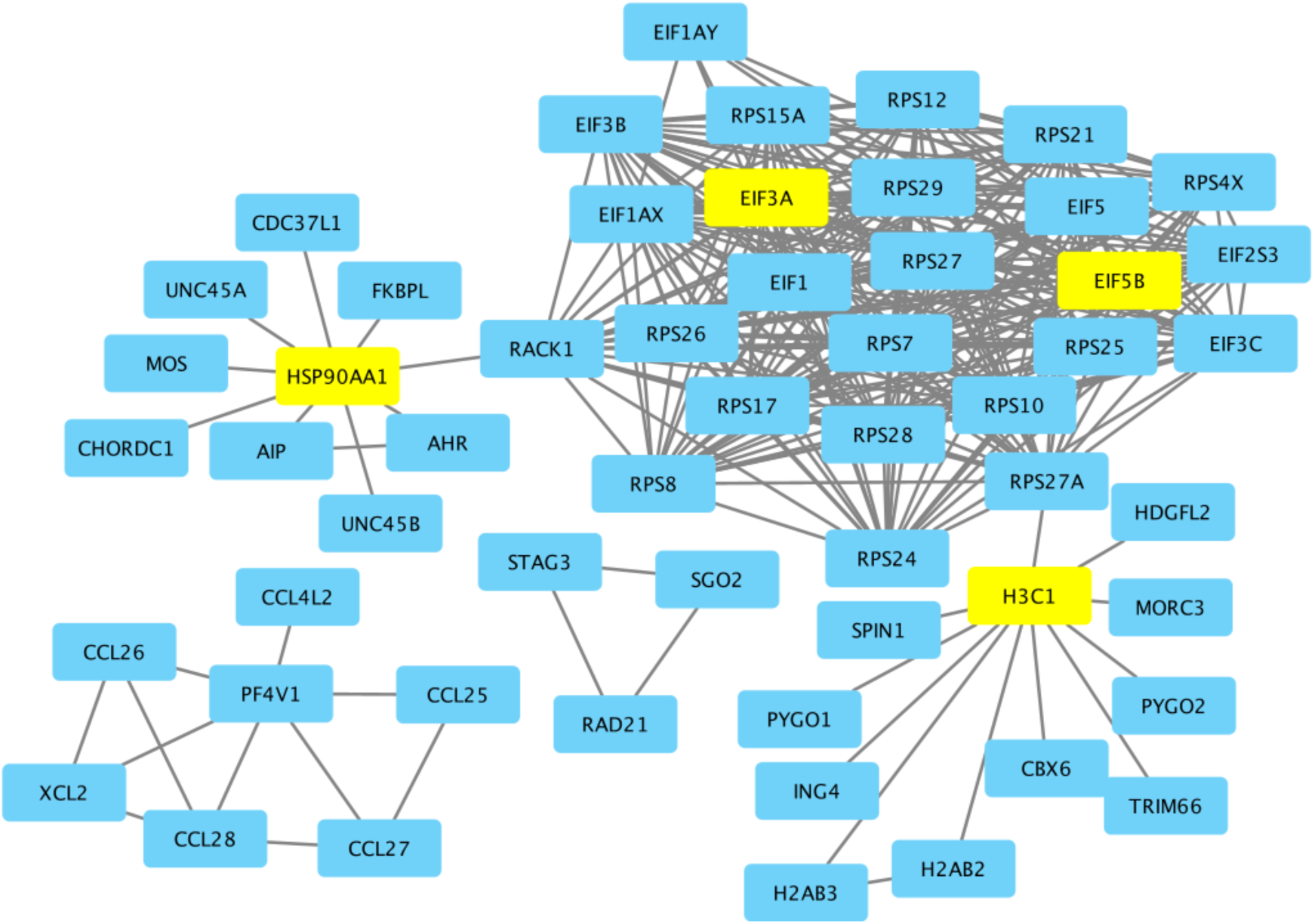
Protein–Protein Interaction Network of Top Genes via STRING. The top 10 protein-coding genes were submitted to STRING. STRING subsequently added intermediary genes to visualize potential interactions using a minimum confidence score of 0.900. Unconnected nodes were removed in Cytoscape. Of the original top genes, only *HSP90AA1*, *EIF3A*, *PF4V1*, *EIF5B*, *H3C1*, and *RAD21* remained in the final network. *HSP90AA1*, *EIF3A*, *EIF5B*, and *H3C1*, are highlighted in yellow as they were the only top genes indirectly connected within the network.

**Table 4.**
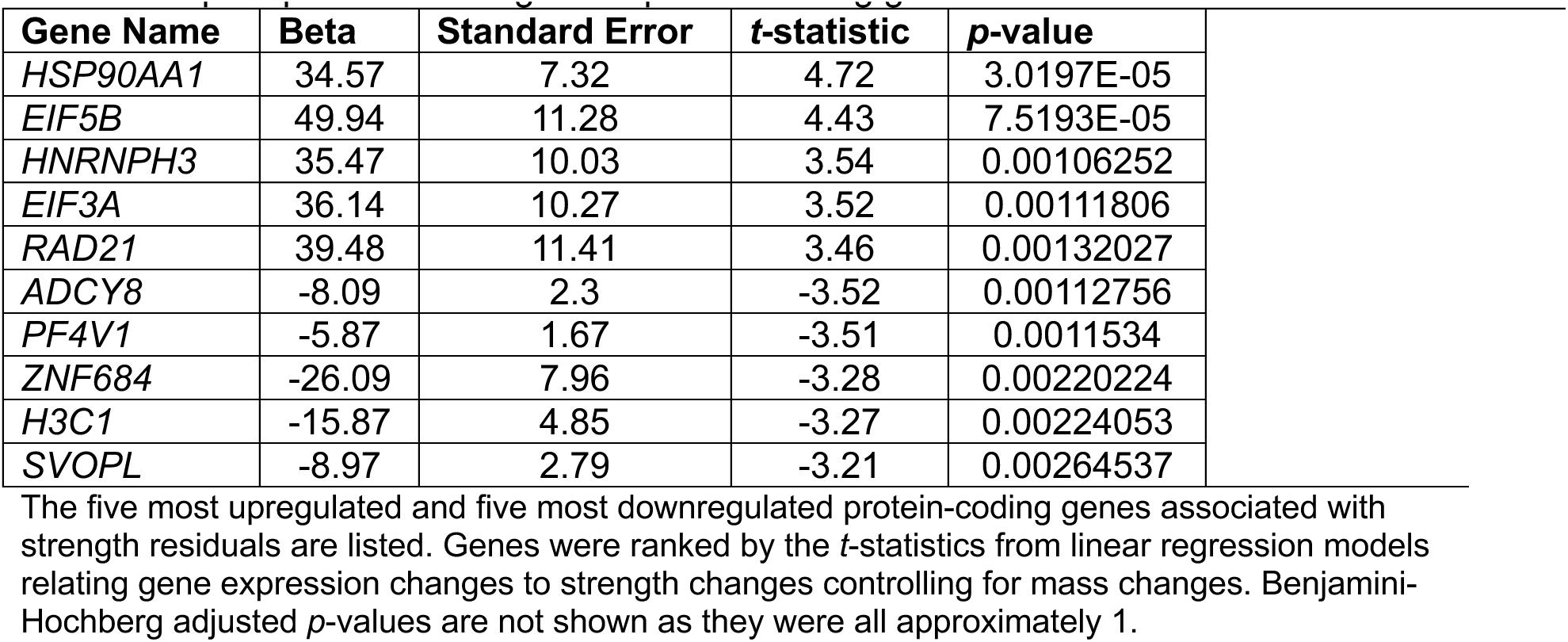
Top 5 up- and downregulated protein-coding genes.

| Gene Name | Beta | Standard Error | $t$-statistic | $p$-value |
| --- | --- | --- | --- | --- |
| *HSP90AA1* | 34.57 | 7.32 | 4.72 | 3.0197E-05 |
| *EIF5B* | 49.94 | 11.28 | 4.43 | 7.5193E-05 |
| *HNRNPH3* | 35.47 | 10.03 | 3.54 | 0.00106252 |
| *EIF3A* | 36.14 | 10.27 | 3.52 | 0.00111806 |
| *RAD21* | 39.48 | 11.41 | 3.46 | 0.00132027 |
| *ADCY8* | -8.09 | 2.3 | -3.52 | 0.00112756 |
| *PF4V1* | -5.87 | 1.67 | -3.51 | 0.0011534 |
| *ZNF684* | -26.09 | 7.96 | -3.28 | 0.00220224 |
| *H3C1* | -15.87 | 4.85 | -3.27 | 0.00224053 |
| *SVOPL* | -8.97 | 2.79 | -3.21 | 0.00264537 |

The five most upregulated and five most downregulated protein-coding genes associated with strength residuals are listed. Genes were ranked by the $t$ -statistics from linear regression models relating gene expression changes to strength changes controlling for mass changes. Benjamini-Hochberg adjusted $p$ -values are not shown as they were all approximately 1.

Finally, GSVA assessed pathway activity at the level of the individual. We were particularly interested in determining changes in pathway activity in individuals with unusually high residual values. Of note, none of the high-residual individuals examined were associated with unusually high or low changes in pathway scores, suggesting these outliers did not provide undue influence on transcriptional findings. We additionally performed correlations between these GSVA change scores and the residuals and found the highest positive correlation coefficients for proliferative pathways and the highest negative correlation coefficients for inflammatory pathways. These results mirrored the GSEA results reported above.

## DISCUSSION

In this secondary analysis of data obtained on healthy, young to middle-aged adults without obesity enrolled in the CALERIE™ trial, we identified gene expression signatures associated with the preservation or enhancement of strength during weight loss over one year. These findings provide new insights into the molecular mechanisms supporting strength maintenance during weight reduction — a vital goal for improving functional outcomes in CR-related geroscience interventions, weight management initiatives, and obesity treatment. As interest in safe and effective CR and obesity treatment strategies continues to grow, understanding how to preserve physical function alongside body weight reduction remains a public health priority.

In our investigation of biological processes supporting strength retention during CR-induced weight loss, GSEA revealed significant enrichment of pathways related to cellular proliferation (MYC targets, E2F targets), immune regulation (TNF-α signaling via NF-κB), and checkpoint control (G2M checkpoint). In addition, the genes *HSP90AA1* and *ADCY8* were identified as the most significant positive and negative *t*-statistic-associated genes respectively. *HSP90AA1* presents itself as an attractive therapeutic target whose modulation could preserve or enhance skeletal muscle strength.^52^ With respect to *ADCY8*, we find these results to be counterintuitive given that cAMP, which is produced by the protein product of *ADCY8*, is known to increase contractile force.^53^ *HSP90AA1* encodes a cytosolic, inducible member of the heat-shock protein 90A (HSP90A) subfamily. HSPs maintain proteome integrity by promoting the correct folding of newly synthesized or denatured proteins, thereby preventing toxic aggregate formation. Emerging evidence supports a key role for HSPs in mediating the beneficial effects of exercise on whole-body and skeletal muscle health. HSP90 proteins can double or triple in concentration in response to stress,^54^ and CR has also been shown to increase cytosolic and nuclear levels of the protein product of *HSP90AA1* – Hsp90-α1.^55^ Hsp90-α1 binds to and promotes the nuclear translocation of the transcription factor TFEB, which promotes autophagy.

Importantly, we previously found CR enhances skeletal muscle quality.^18^ *HSP90AA1* was among the genes upregulated by CR in skeletal muscle. Collectively, these findings suggest *HSP90AA1* could enhance skeletal muscle quality through cytosolic and mitochondrial proteome maintenance and autophagy.^56^ Importantly, multiple lines of evidence support the role of *HSP90AA1* in maintaining skeletal muscle health and function. For example, overexpression of mouse *hsp90aa1* was associated with increased expression of a range of myosin heavy chain genes, an increase in cytosolic myosin, and an increase in the myosin replacement rate.^57^ *Hsp90aa1* has also been shown to be essential for myofibril organization in skeletal muscles of zebrafish embryos.^58^ Targeting *HSP90AA1* may therefore represent a strategy to preserve muscle strength during weight reduction via enhanced protein folding and autophagy. Future studies should investigate *HSP90AA1*’s functional role through knockout, knockdown, or overexpression models to directly test its contribution to strength maintenance or enhancement.

CR also enhances satellite cell proliferation in both mice and humans.^59,60^ Our current GSEA findings – implying enrichment of pathways linked to cellular proliferation and muscle adaptation (MYC Targets, E2F Targets, G2M Checkpoint) – strongly suggest strength gains may involve enhanced proliferation of muscle progenitor cells. Alternatively, increased proliferation of other muscular cell types, like myocytes, may contribute to this signal. Although hypertrophy is typically regarded as the primary driver of strength, our results suggest hyperplasia may also play a role and merits further investigation.

We also observed a positive association between increased protein secretion and strength. In earlier studies, we found CR markedly enhances hepatic protein secretion.^59^ Although no change in skeletal muscle protein secretion was detected, the analysis had limited sensitivity for low molecular weight (MW) proteins. Skeletal muscle secretes a class of small MW proteins known as myokines, which promote functional improvements across multiple organs.^61^ Thus, the protein secretion signature associated with greater strength may reflect increased myokine secretion.

Further, we identified strong associations between metabolic pathway alterations and strength. Overall, our findings suggest a shift towards oxidative phosphorylation is related to strength gains. In contrast, several inflammatory pathways, including IL6-JAK-STAT3 signaling, were negatively associated with strength. These results are consistent with prior evidence indicating systemic IL6 contributes to declines in skeletal muscle strength.^62^ Together, these findings highlight several pathways that could be targeted to preserve or enhance strength, even during mass loss.

Together, these mechanistic insights have important translational implications for obesity treatment and weight management. As pharmacologic and surgical interventions for obesity become increasingly common,^63,64^ evaluating the roles of these genes in the context of drug- or surgery-induced weight loss could clarify their relevance across diverse treatment strategies. Such studies may inform the development of targeted interventions to preserve muscle strength during rapid weight reduction, thereby maintaining muscle function and quality – both essential for long-term health and quality of life.

A major motivation for this study was to identify genes and pathways whose modulation could preserve or even enhance skeletal muscle strength in the context of CR-induced weight loss. This is particularly relevant given the growing use of glucagon-like peptide-1 receptor agonists (GLP-1 RAs) for obesity treatment, which primarily act by reducing caloric intake through appetite suppression. A common, potentially adverse, side-effect of these agents is substantial loss of lean body mass.^65^ To address this side-effect, several studies have tested t co-administered agents that preserve or even increase lean mass during GLP-1 RA-induced weight loss. For example, Mastaitis et al. demonstrated that the dual blockade of GDF8 and activin protects against GLP-1-induced muscle loss in mice and non-human primates.^66^ Despite these promising results, we question the long-term effectiveness of promoting growth pathways during CR and weight loss. In contrast, our analytical approach identifies genes directly related to strength independent of whole-body mass, thereby prioritizing pathways that may sustain muscle function without promoting excessive anabolic signaling.

This goal is particularly important for older adults, who are at greater risk of sarcopenia, frailty, and mobility impairments.^67^ The “obesity paradox” further underscores the delicate nature of weight loss during older adulthood.^68,69^ Although the present study focused on younger and middle-aged adults without obesity, the molecular pathways identified here may inform future efforts to promote strength preservation in older adults undergoing weight loss, ultimately supporting better functional outcomes and quality of life.

This secondary analysis is not without limitations. We focused on a select subset of genes – those most strongly correlated with strength retention – to generate interpretable results and demonstrate the feasibility of this analytical pipeline. A more comprehensive evaluation of all significant genes could provide additional insights into biological pathways contributing to strength retention during weight loss. Additionally, transcriptomic data were available for a relatively small number of participants at both timepoints. Future studies should therefore pursue broader pathway analyses with experimental validation. Our linear regression model assumed a constant linear relationship between changes in whole-body mass and strength; however, this relationship may be nonlinear, with greater weight losses disproportionately reducing strength. Future work should explore nonlinear modeling approaches to better capture the biological complexity.

## Conclusions

In summary, this analysis identified several molecular pathways and candidate genes associated with strength preservation during CR-induced weight loss in healthy adults. Among those identified, *HSP90AA1* emerged as a particularly compelling target due to its established biological relevance and potential role in maintaining skeletal muscle quality. Future research should examine the functional contribution of *HSP90AA1* to strength preservation using both animal models and larger human cohorts, particularly in combination with exercise-based interventions commonly used to mitigate muscle loss during weight reduction. A deeper understanding of the molecular mechanisms governing muscle preservation and strength enhancement may ultimately support the development of targeted, molecular-based strategies to maintain muscle quality and functional strength during weight loss and obesity treatment, thereby improving long-term health outcomes.

## Acknowledgments

The authors would like to thank the CALERIE™ participants, as well as staff instrumental in conducting the baseline and 12-month assessments and the intervention.

---

**Supplementary Figure 1.**
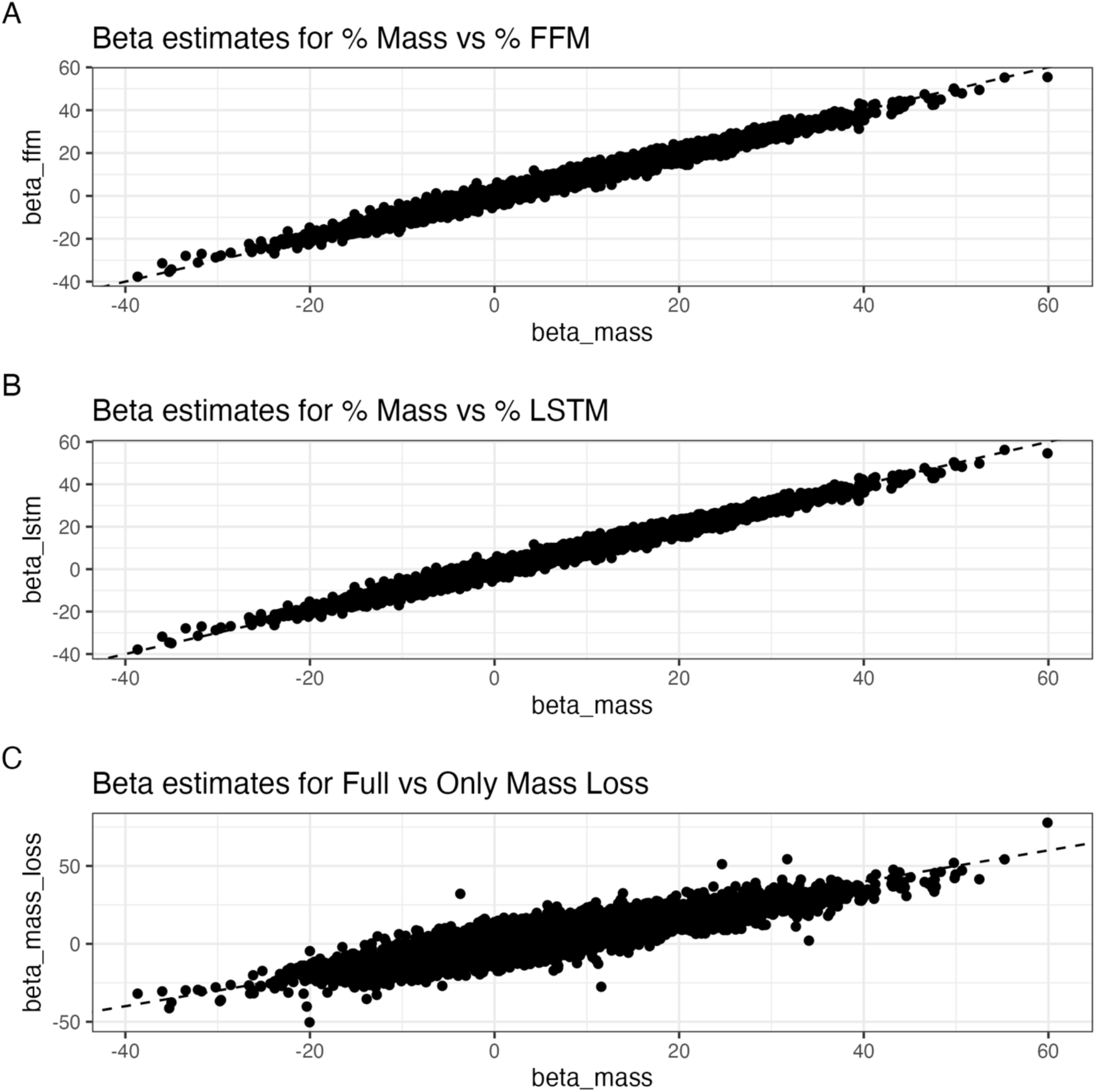
Multiple regression sensitivity analyses. Comparison of gene-associated beta coefficients for models controlling for change in whole-body mass compared to models controlling for change in fat-free mass (ffm) (A); or compared to models controlling for change in lean soft-tissue mass (lstm) (B); or compared to models that only used data from subjects who had lost whole-body mass (N=33) (C).

